# Insects visit *Fusarium xyrophilum* pseudoflowers on the host *Xyris surinamensis* (Xyridaceae) and carry fungal DNA on their bodies

**DOI:** 10.1101/2024.03.05.583517

**Authors:** Terry J. Torres-Cruz, Tristan M. Cofer, Laura M. Kaminsky, Lauren A. Ré, Jack R. Johnson, Alexis Vaughn, James H. Tumlinson, Imane Laraba, Robert H. Proctor, Hye-Seon Kim, Terrence H. Bell, M. Catherine Aime, Michael J. Skvarla, David M. Geiser

**Author notes:** Corresponding author: David M. Geiser.

## Abstract

The fungus *Fusarium xyrophilum* produces flower-like structures (i.e., pseudoflowers) that were recently discovered on yellow-eyed grasses (*Xyris* spp.) in Guyana. It is unknown whether these pseudoflowers, which are composed entirely of fungal tissue, are true mimics that attract insects as a means of fungal dispersal. We evaluated the potential of *F. xyrophilum* to affect insect visitation patterns to flowers and pseudoflowers by 1) documenting insect visitation to *X. surinamensis* in Guyana, 2) measuring the presence of *F. xyrophilum* DNA on insects, and 3) evaluating fluorescence and volatile production on flowers and pseudoflowers. We report for the first time Vespidae, Formicidae, Salticidae, Acrididae, and Tetrigidae visiting *Xyris*. Diverse insects, including Conocephalini spp. (meadow katydids; Tettigoniidae), *Camponotus* spp. (carpenter ants; Formicidae), and a Geometridae sp. (geometer moths) were found to visit flowers and pseudoflowers. *Fusarium xyrophilum* DNA was detected on 3/12 (25%) of captured insect bodies using conventional and quantitative PCR. Volatiles produced in the field by pseudoflowers and flowers were similar, except for the presence of a sesquiterpene, putatively identified here as α-gurjunene, which was detected both in *F. xyrophilum* pure cultures and field-collected pseudoflower samples, but not from flowers. The production of this sesquiterpene by *F. xyrophilum* and the fluorescence of *X. surinamensis* peduncles represent potential signals involved in insect attraction for this system. These observations, along with the overlap in insect visitors of flowers and pseudoflowers and the detection of *F. xyrophilum* DNA on insect bodies, are consistent with insect visitors being vectors of *Xyris* pollen and *F. xyrophilum* propagules between host plants.

## INTRODUCTION

*Fusarium* (Nectriaceae) is a ubiquitous genus of filamentous ascomycetes. The genus includes many plant pathogens of agronomic importance, opportunistic human pathogens, and mycotoxin producers (O’Donnell et al. 2013; Ma et al. 2013). Approximately 80% of all cultivated plants have at least one disease attributed to a *Fusarium* species; including head blights/ear rots on cereals and vascular wilts and root rots on vegetables and several other crops (Kvas et al. 2009). Exploration of new ecosystems continues to reveal novel *Fusarium* diversity with varied arrays of ecological functions and potential applications.

A recently described species, *F. xyrophilum*, produces flower-like structures (i.e., pseudoflowers) on species of *Xyris* (yellow-eyed grass; Poales: Xyridaceae). *Fusarium xyrophilum* is a member of the *F. fujikuroi* species complex (FFSC), one of the most studied *Fusarium* species complexes that comprises diverse mycotoxigenic and plant pathogenic species (Kvas et al. 2009). Some other FFSC species induce morphological changes to plant inflorescences (Marasas et al. 2006; Freeman et al. 2014), but *F. xyrophilum* is the only *Fusarium* species reported to form pseudoflowers (Laraba et al. 2020a, 2020b). The pseudoflowers are yellow-orange discoid/lobulate masses that emerge at the tip of the inflorescence. These pseudoflowers can last for several days (Laraba et al. 2020a), unlike the bright yellow *Xyris* flowers, which emerge at a rate of one or two per day and open for only a few hours from mid-morning to mid-afternoon before shriveling (Kral 1988). The first report of pseudoflowers on *Xyris* dates to 1988, when orange masses noted on spikes of *X. subglabrata* and *X. setigera* were attributed to a “smut fungus” infecting the flowers. Despite these reports, no further research on these structures was published until 2020, when pseudoflowers were noted on *X. surinamensis*, *X. setigera* and *X. bicephala* (Laraba et al. 2020b). Furthermore, a survey of specimens in three United States herbaria revealed the presence of pseudoflowers preserved on accessions of *X. surinamensis*, *X. setigera*, and *X. subglabrata*, including some specimens archived since 1919 (Laraba et al. 2020b). The close visual resemblance of *F. xyrophilum* pseudoflowers to *Xyris* flowers suggests a newly discovered fungal-plant mimicry system (Laraba et al. 2020a, b).

Pseudoflowers are thought to increase insect visitation to plants through color and scents (Roy 1994; Naef et al. 2002; McArt et al. 2016) to facilitate diverse ecological functions like fertilization of fungi during sexual reproduction, dissemination of spores, or faciliting entry of fungi into the host without eliciting a defense response (Ngugi and Scherm 2006). The attraction of insects to plant hosts by pseudoflower-inducing fungi has been studied in a handful of mimicry systems. For example, in mummy berry disease caused by *Monilinia vaccinii-corymbosi* (*Mvc*), infected blueberry leaves reflect ultraviolet (UV) light, providing a visual signal to pollinators that is similar to that of healthy flowers (Batra and Batra 1985). Infected leaves also mimic the floral scent of blueberry due to the presence of the bee-attracting volatiles cinnamyl alcohol and cinnamic aldehyde in the leaves (McArt et al. 2016). Several insects, including bees and flies, are attracted to the infected flowers and leaves and carry *M. vaccinii-corymbosi* conidia on their bodies likely acting as fungal vectors. However, not all fungus-induced pseudoflowers visually mimic uninfected flowers of their plant hosts. Pseudoflowers induced by *Puccinia monoica* on *Boechera stricta* (rockcress) mimic other plants in the vicinity that bloom at the same time, such as *Ranunculus inamoenus* (buttercups). These pseudoflowers emit a fragrance consisting mostly of aromatic alcohols, aldehydes, and esters, while *B. stricta* flower scent is a blend of terpenoids and aliphatic green leaf volatiles (Raguso and Roy 1998). A similar trend is observed for *P. arrhenatheri* pseudoflowers and *Berberis vulgaris* flowers, which share two volatiles (Naef et al. 2002). In these cases, the pseudoflowers do not chemically mimic floral scents, but mimic them functionally, as both attract pollinators. When comparing single-species plots to mixed plots of buttercups and infected rockcress, both buttercups and pseudoflowers receive more visits when they are both present in the same plot than when alone (Roy 1994). The difference in the production of volatiles between pseudoflowers and host flowers may account for this increased pattern of insect visitation in mixed plots because the emission of a diverse array of compounds attracts more diverse insects (Raguso and Roy 1998).

Most knowledge of insect visitation to *Xyris* has been carried out in temperate regions, where *Xyris* species were hypothesized to be wind-pollinated due to the lack of nectaries, and ‘infrequent’ visitation by pollen-collecting andrenid bees (Kral 1983). However, *X. tennesseensis* is visited during anthesis mostly by halictid bees (Hymenoptera: Halictidae) and pollen-consuming syrphid flies (Diptera: Syrphidae) (Boyd et al. 2011; Moffett and Boyd 2013). *Lasioglossum zephyrus* (Hymenoptera: Halictidae) manipulates *X. tennesseensis* flowers to open prematurely, ensuring first access to floral rewards (Wall et al. 2002). Observations of Halictidae and Syrphidae are also reported for *X. asperulla* and *X. tortulla* (Freitas and Sazima 2006). Seed heads of *X. iridifolia* are presumed to be larval food for the moth *Coleophora xyridella* (Lepidoptera: Coleophoridae), which produces clusters of cigar-shaped, tan-colored coleophorid cases that attach to seed heads (Landry 2005). A recent study provided the first documentation of arthropods in the orders Araneae, Coleoptera, and Orthoptera on *Xyris* spp. in Guyana that are likely to carry pollen on their bodies and feed on pollen and petals, suggesting that arthropods could play a role in *Xyris* pollination (Torres-Cruz et al. 2024).

Limited information is available on the extent to which the presence of *F. xyrophilum* pseudoflowers affect the interactions with *Xyris* insect visitors and the participation of insects in dispersing this fungus. Therefore, our objectives for this study were to **1)** assess insect visitation to *X. surinamensis* flowers and pseudoflowers in Guyana; **2)** determine if *F. xyrophilum* is detected molecularly on insects; and **3)** compare fluorescence and emission of volatile organic compounds (VOCs) produced by pseudoflowers and flowers of *X. surinamensis* in the field. Our study provides the first studies of insect visitation and VOCs production on *X. surinamensis* bearing true flowers and *F. xyrophilum* pseudoflowers. Along with the discovery of *F. xyrophilum* DNA on insect bodies confirming the hypothesis that insects vector *F. xyrophilum*, we have largely expanded the current knowledge of this recently discovered mimicry system.

## METHODS AND MATERIALS

### Study Area

Observations and sample collection were conducted in Guyana between 15 December 2021 and 2 January 2022. Two sites (1 and 2) were studied in the Demerara-Mahaica Region. The vegetation at both sites was comprised of short flowering trees (e.g., *Clusia* sp., Malpighiaceae), sedges, rushes, and grasses, including a variety of *Xyris* species. (e.g., *X. surinamensis*, *X. involucrata*). The surface of the white sand soil between plants was fairly covered by a diversity of mosses (e.g., *Sphagnum*), as well as *Drosera kaeiteurensis*, *D. intermedia*, and Eriocaulaceae. Diverse plants with purple flowers (e.g., *Burmannia bicolor*, *Chelonanthus purpurascens*, *Sauvagesia* sp., multiple species of Melastomataceae) and yellow flowers (e.g., *Xyris* spp., *Perama hirsuta*, *Utricularia juncea*, *Chamaecrista* sp.) are found in the area. During sampling days, Site 1 (6° 26’ 40’’ N 58° 11’ 27’’ W; ∼65 m.a.s.l.) showed temperatures ranging from 22–38° C; while in Site 2 (6° 19’ 10’’ N 58° 12’ 13’’ W; ∼60 m.a.s.l.) temperatures varied from 28–35° C. Photography of plant, insect, and fungal observations are available in iNaturalist under the project: “<u>Ecosystem profiles of *Xyris* Research Sites</u>”. The incidence of plants bearing pseudoflowers was recorded by counting the number of dead and living *Xyris* plants with and without pseudoflowers in five 1 × 30 m nonoverlapping transects at the two sites in Demerara-Mahaica region. Data collection took place on 1 January 2022 at Site 1 and 2 January 2022 at Site 2.

### Insect Collection from *Xyris surinamensis*

Insects were captured at both collection sites (1 and 2) in Demerara-Mahaica Region using yellow pan traps to facilitate insect identification in areas with *X. surinamensis*. Twelve-ounce yellow plastic bowls (Party Solids, Kingston, PA) were filled with ∼200 ml of water mixed with two drops of unscented biodegradable dish soap as a surfactant. Three bowls were placed in a random pattern at ground level within 0.5–1 m of *X. surinamensis* plants for approximately 6 hours each day (9:00-15:00) over a period of seven days, totaling six bowls at Site 2 (two days of collection) and 15 bowls at site 1(five days of collection). Differences in sampling efforts (two vs five days of collection) depended on the availability of opened flowers at each collection site on certain dates. At the end of the day, insects collected in each bowl were transferred into vials filled with 90% ethanol. Arthropods were pinned and identified using morphology.

### Arthropod Visitation to *Xyris surinamensis*

Time-lapse video was used to assess arthropod visits to *X. surinamensis* plants bearing flowers in comparison to plants bearing *F. xyrophilum* pseudoflowers at sites 1 and 2 (Demerara-Mahaica Region). Video clips were recorded between 9:00 and 16:00, based on patterns of insect visitation. Time-lapse pictures with 5 s intervals were captured using two GoPro Hero7 Black cameras (GoPro Inc., San Mateo, USA) attached to extended batteries (refuel RF-6H50; Mizco International, Avenel, New Jersey) to provide longer battery life throughout the day. At each site, one camera was focused on a spike bearing a pseudoflower while the other on a true flower-bearing spike. Cameras were positioned ∼15 cm away from their respective targets, using tripods for stability, and were provided with a macro lens (52 mm, 10x magnification). Approximately 40 min of recording was obtained at each target before moving the camera to a new flower or pseudoflower at the same collection site. Approximately seven ∼40 min recordings per target type (i.e., pseudoflower or flower) were taken per field day. A total of 78 hours 13 min and 25 s of captured video footage was obtained over 10 days of fieldwork. Video data were assessed individually in duplicate. Morphological identity of the insect (i.e., order, family, or genus), contact with the target, and interaction time with the specific target were obtained for each observation. The number of insect visitors, type of *Xyris* tissue visited and types of insect visitors were compared between *Xyris* flowers and *F. xyrophilum* pseudoflowers. Video clips captured herein are available on YouTube as playlist “*<u>Fusarium xyrophilum</u>* <u>– *Xyris surinamensis* insect</u> <u>visitation study</u>” (playlist ID: PL19pSjmfC9cSceih6nlMnaBQj57ITfvq8).

### Detection of *F. xyrophilum* on Arthropod Visitors

Due to the size and fragility of *Xyris* flowers and the frequency of insect visitation observed, hand-netting was not feasible. Efforts were made to hand-collect insects using tweezers. Two researchers canvased the survey area at Site 1 in Demerara-Mahaica Region for about an hour on each sampling date to hand-collect insects in contact with plants bearing pseudoflowers or true flowers. Insects collected (n=12) were preserved individually in 90% ethanol to prevent cross-contamination. Genomic DNA was extracted from the full body of insects using the DNeasy Plant MiniKit (Qiagen) following the manufacturer’s instructions. To assess the presence of *F. xyrophilum* on insect bodies, we PCR-amplified a ∼173-bp fragment of the ribosomal intergenic spacer region (IGS rDNA) using primers IGS-1f and IGS-1r (Laraba et al. 2020b). DNA extracted from pure *F. xyrophilum* cultures was used as a positive control and sterile ultrapure water was used as negative control in the PCR reactions. Additionally, insects were identified by amplifying a ∼636 bp portion of the mitochondrial cytochrome c oxidase subunit I (COI) using the primer pair LCO1490 and HC02198 (Folmer et al. 1994). IGS rDNA and COI amplicons were Sanger sequenced at the Penn State Genomics Core Facility (University Park, PA). Sequences were submitted to GenBank under accession numbers OQ121925–27 and OQ152379–90, respectively.

### qPCR Quantification of *F. xyrophilum* Using IGS rDNA

To determine the amount of *F. xyrophilum* biomass on arthropods visiting the pseudoflowers, a qPCR was performed using the same IGS rDNA primers used for conventional PCR above. Each 20 μL reaction contained 10 μL of 2X QuantiNova SYBR Green Master Mix (Qiagen, Redwood City, CA, USA), 1.4 μL of each primer (10 μM), 6.2 μL of sterile nuclease-free water, and 1 μL of template DNA. Negative controls contained 1 μL of sterile nuclease-free water instead of template DNA and were included on every run. All qPCR reactions had three technical replicates and were carried out using a Bio-Rad C100 Touch Thermal Cycler and CFX96 Real-Time System machine (Bio-Rad, Hercules, CA, USA). All reactions were run using the manufacturer’s recommended cycling conditions for the QuantiNova SYBR Green PCR kit. Melting curve analysis consisted of 5 s at every 0.5 °C interval from 65 °C to 95 °C. The DNA concentration of a cleaned amplicon of *F. xyrophilum* was measured using a Qubit 3 fluorometer and 1X dsDNA High Sensitivity Assay Kit (Invitrogen, Waltham, MA, USA) and the number of amplicon molecules per μL was calculated. This was then diluted to a stock solution of 3.092 × 10^-6^ molecules per μL as the highest concentration standard (standard 1). Seven additional standards were prepared via 8-fold standard dilutions. Each standard was run in triplicate using the qPCR cycling conditions and reaction volumes detailed above. The slope of the resulting calibration curve of quantification cycle (Cq) values vs. log10 of amplicon molecules initially present in the PCR tube was used to calculate assay efficiency according to the following equation: PCR efficiency = 10^-1/slope^ – 1 (Bustin et al., 2009). To determine if there was qPCR inhibition caused by the extracted DNA sample, we followed a method similar to (Kaminsky and Bell 2022). Five concentrations of insect DNA were prepared from the 12 insect extractions obtained in the previous step: undiluted, 5-fold, 10-fold, 50-fold, and 100-fold dilutions in sterile nuclease-free water. qPCR reactions were conducted, as described before, with template consisting of 1 μL of diluted insect DNA plus 1 μL of either standard 1 (3.092 × 10^6^ molecules per μL), standard 3 (4.831 × 10^4^ molecules per μL) or standard 5 (7.549 × 10^2^ molecules per μL), and 1 μL less of sterile water to maintain a 20 μL reaction volume.

The Cq values of each insect dilution-standard combination were compared to that of the corresponding standard run alone. An increase in Cq value in the presence of insect DNA was not observed, indicating no qPCR inhibition. Subsequently, the qPCR reactions in this experiment were conducted using undiluted insect DNA. Cq values and melting curve results were determined with the Bio-Rad CFX Maestro software using default settings. A subset of qPCR amplicons was sequenced at the Penn State Genomics Core Facility (University Park, PA), as previously detailed, to confirm their identity as *F. xyrophilum*.

### Volatile Organic Compounds Produced by *X*

surinamensis *Bearing Flowers vs. Pseudoflowers.* Volatile organic compounds (VOCs) were collected from the headspace of randomly selected *F. xyrophilum-*infected *X. surinamensis* plants bearing pseudoflowers and *X. surinamensis* plants bearing true flowers at Sites 1 and 2 in the Demerara-Mahaica Region. Volatiles were collected between 10:00 am and 4:00 pm on four different days by placing bouquets of seven flowers or seven pseudoflowers into 8-oz glass chambers placed in the field. Bouquets were prepared by excising inflorescences of *X. surinamensis* a few centimeters below the spike and wrapping the stems in wet paper towels covered with aluminum foil. Each bouquet was placed into a glass chamber and closed with a lid fitted with two SuperQ filters (20 mg; Alltech Associates, Deerfield, IL, USA), one to clean the air coming into the chamber and the other to trap VOCs released in the headspace. Air was pulled from chambers at ∼1 L min^-1^ using a 9V battery-powered vacuum pump for 1.5 – 3 hours. In addition to collecting VOCs emitted from flowers and pseudoflowers, at each run a control glass chamber was set up without any samples to account for background corrections. A total of seven collection runs were conducted for each sample type, including the control. Filters were transported to the lab, eluted with 100 µL of n-hexane/dichloromethane (1:1, v/v) containing 500 ng of nonyl acetate as an internal standard and analyzed by gas chromatography-mass spectrometry (GC-MS).

To gain better insight into the VOCs profile of *F. xyrophilum*, VOCs were collected from pure cultures of *F. xyrophilum* (NRRL 62721, FRC M-8921), *F. verticillioides* (NRRL 20956, FRC M-3125), and *F. thapsinum* (NRRL 22048, FRC M-6562). The two latter species, genetically closely related to *F. xyrophilum* as part of the African clade of FFSC, were used to determine olfactory signatures specific to *F. xyrophilum*. The strains were cultured in duplicate on Potato Dextrose Agar (PDA) in 16-oz glass chambers covered with aluminum foil and incubated at room temperature under 12h/12h light/dark. After three weeks, VOCs were collected in duplicate for 30 min using a 100 µm polydimethylsiloxane solid-phase micro extraction (SPME) fiber (Supelco, Bellefonte, PA) that was introduced inside the glass chamber and exposed to the headspace above the fungus. Sterile PDA jars were used as control. Samples were analyzed on an Agilent 6890 series gas chromatograph (GC) coupled to an Agilent 5973 quadrupole mass selective detector (MS; interface temperature, 250 °C; quadrupole temperature, 150 °C; source temperature, 230 °C; electron energy, 70 eV). Samples were injected in splitless mode onto an HP-5MS column (30 m × 0.25 mm i.d. × 0.25 µm thickness; Agilent, Palo Alto, CA, USA) using helium as the carrier gas at a constant flow rate of 1 mL min^-1^. For liquid samples, the oven temperature was held at 40 °C for 2 min, then increased from 40 to 100 °C at 8 °C min^-1^, 100 to 160 °C at 5 °C min^-1^, 160 to 260 °C at 40 °C min^-1^, and held at 260 °C for 7 min. For SPME samples, the oven temperature was held at 40 °C for 2 min, then increased from 40 to 160 °C at 4 °C min^-1^, 160 to 280 °C at 30 °C min^-1^, and held at 280 °C for 4 min. Compounds were identified by comparing the mass spectrum to those in the NIST14 library.

Three previously generated *F. xyrophilum* genome sequences (GenBank accessions VYXA00000000, VYWZ00000000, and VYWY00000000; Laraba et al. 2019) were examined using two methods to identify putative terpene synthase genes potentially involved in production of terpene VOCs (see results). First, the genome sequences were subjected to antiSMASH version 4 analysis following the same approach detailed in Kim et al. (2020). Second, a database of predicted proteins from the three *F. xyrophilum* genomes were subjected to BLASTp analysis using query sequences consisting of 40 coding region sequences representing the breadth of phylogenetic diversity of a previously analyzed set of terpene synthase genes from filamentous fungi (Agger et al., 2009) and all terpene synthase genes described in the species *Fusarium fujikuroi* (Niehaus et al., 2016).

*UV Fluorescence. Xyris surinamensis* flowers and *F. xyrophilum* pseudoflowers from Site 1 in the Demerara-Mahaica region were exposed to ultraviolet light generated by a Convoy C8 365nm UV light (Yooperlite, Brimley, MI) to evaluate their UV reflection. Photographs of flowers were taken in the field using an Olympus Tough TG-6 camera. Additionally, three flowers and pseudoflowers were photographed in the dark approximately 6 hours after being collected from the field using a Sony a7riii with a Sigma 105mm 2.8 DG DN macro lens.

## RESULTS

### Arthropods in the vicinity o*f Xyris surinamensis*

A total of 204 insects representing six orders and at least 25 families were collected in yellow pan traps (**Table 1**), along with 3 arachnids from two sites of *X*. *surinamensis* populations (Site 1 and 2) in the Demerara-Mahaica Region of Guyana. Additionally, three of the captured insect specimens were not identifiable to order level (XYR006, XYR056, XYR059). Diptera and Hymenoptera were the most collected insects at the collection sites, with 111 and 65 individuals, respectively. The most represented families were Dolichopodidae (74 specimens) and Formicidae (53 specimens). A comprehensive list of the 204 arthropods found in the vicinity of *X. surinamensis* populations at Demerara-Mahaica, Guyana with their genus and species can be found in **Supplementary Material 1**.

**Table 1.**
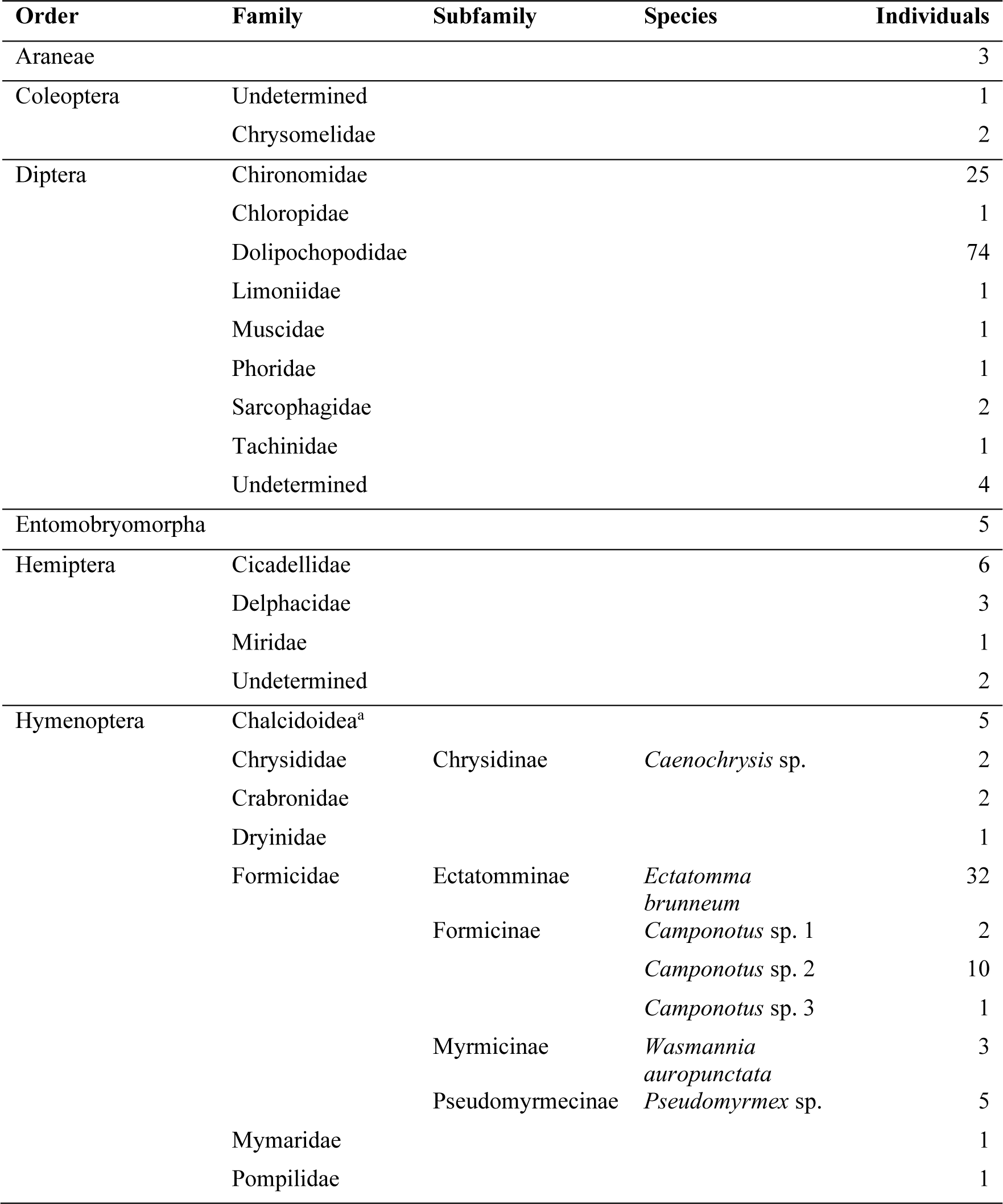
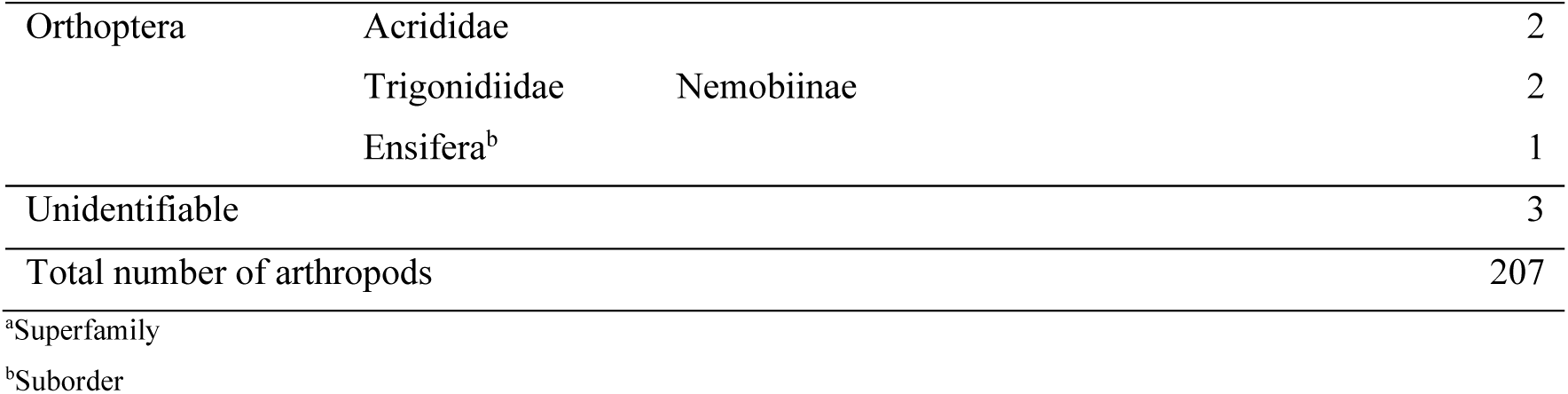
Arthropods in the vicinity of *Xyris surinamensis* in the Demerara-Mahaica region of Guyana.

### Arthropod visitation to Xyris surinamensis

In 10 days of field observation, a total of 78 h 13 min and 25 s of video footage were captured as timelapse to document insect visitation to *X. surinamensis* plants. Of these, 34 h 46 min 30 s were recorded to study insects visiting *X. surinamensis* flowers and 43 h 26 min 55 s for pseudoflowers. Over 10 days at Site 1, we found that on average >10 flowers opened per day, with only two collection dates where <10 flowers were open with an average of 1.53% incidence of pseudoflowers (**Table 2**). In contrast, the number of opened flowers at Site 2 varied across collection dates from zero opened flowers to <5 in a day. Thus, more video data were collected for pseudoflowers than flowers at this site. The incidence of pseudoflowers at Site 2 was on average 4.75%. Even with this data collection bias, fewer insect visitors were observed in contact with pseudoflowers (n = 4) than flowers (n = 19; **Table 3**).

**Table 2.** Percent incidence of pseudoflowers on *Xyris surinamensis* in Demerara Mahaica, Guyana based on transect counts.

| Transect | Site 1 |  |  | Site 2 |  |  |
| --- | --- | --- | --- | --- | --- | --- |
|  | *Xyris* without pseudoflowers | *Xyris* with pseudoflowers | % incidence | *Xyris* without pseudoflowers | *Xyris* with pseudoflowers | % incidence |
| 1 | 267 | 6 | 2.20 | 1072 | 56 | 4.96 |
| 2 | 267 | 14 | 4.98 | 570 | 29 | 4.84 |
| 3 | 444 | 0 | 0.00 | 370 | 8 | 2.12 |
| 4 | 543 | 7 | 1.27 | 70 | 6 | 7.89 |
| 5 | 606 | 6 | 0.98 | 103 | 10 | 8.85 |
| **Overall** | **2127** | **33** | **1.53** | **2185** | **109** | **4.75** |

**Table 3.**
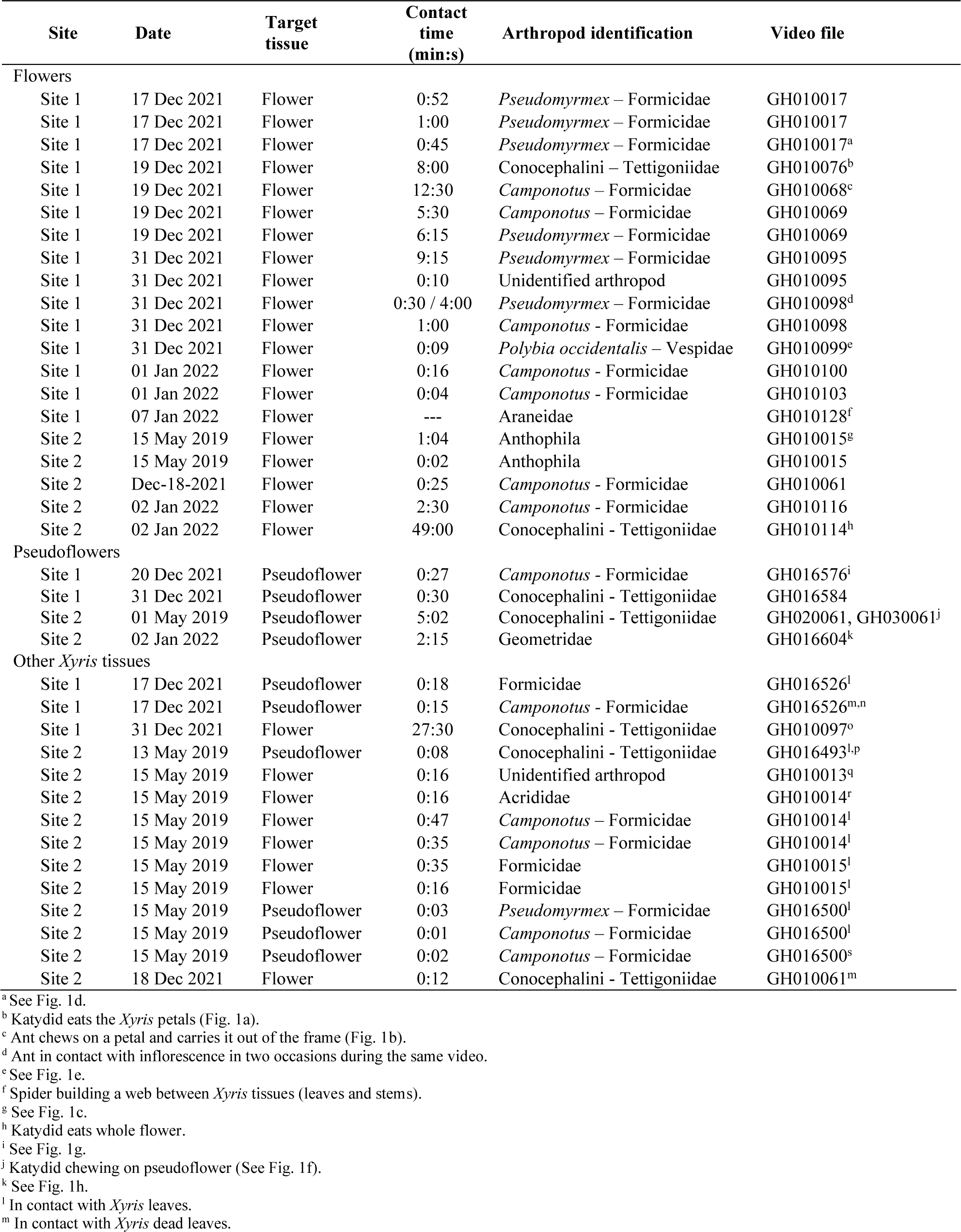
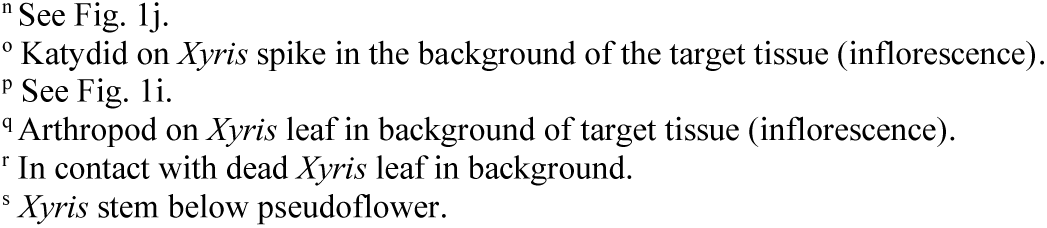
Arthropod visitors of *Xyris surinamensis* flowers, *Fusarium xyrophilum* pseudoflowers, and other tissues.

The same insects were observed to be in contact with pseudoflowers and flowers (**Table 3**). This includes two observations of Conocephalini spp. found chewing on flowers (*e.g.,* **Figs. 1a, n**) and two observed feeding on pseudoflowers (*e.g.,* **Fig. 1f**); seven observations of *Camponotus* spp. on *Xyris* spikes bearing a flower (*e.g.,* **Fig. 1d**), including one observed carrying a petal out of the frame (**Fig. 1b**), and one *Camponotus* spp. on a pseudoflower (**Fig. 1g**); and one Geometridae species attached to pseudoflower tissue (**Fig. 1h**) as well as below the spike bearing a flower (**Fig. 1m**). However, it is worth nothing that some arthropods were exclusively in contact with flowers but not pseudoflowers (**Table 3**). These include two Anthophila species (*e.g.,* **Fig. 1c**), a Vespidae sp. (**Fig. 1e**), an Araneidae sp., and seven observations of Formicidae spp. (*e.g.,* **Fig. 1d**). Additionally, one Salticidae sp. was found in contact with the spike of *X. surinamensis* below the pseudoflower structure (**Fig. 1k**) and on a couple of occasions spider webs were observed between *Xyris* tissues (*e.g.,* scapes, spikes) and the pseudoflowers (**Fig. 1l**).

**FIG. 1.**
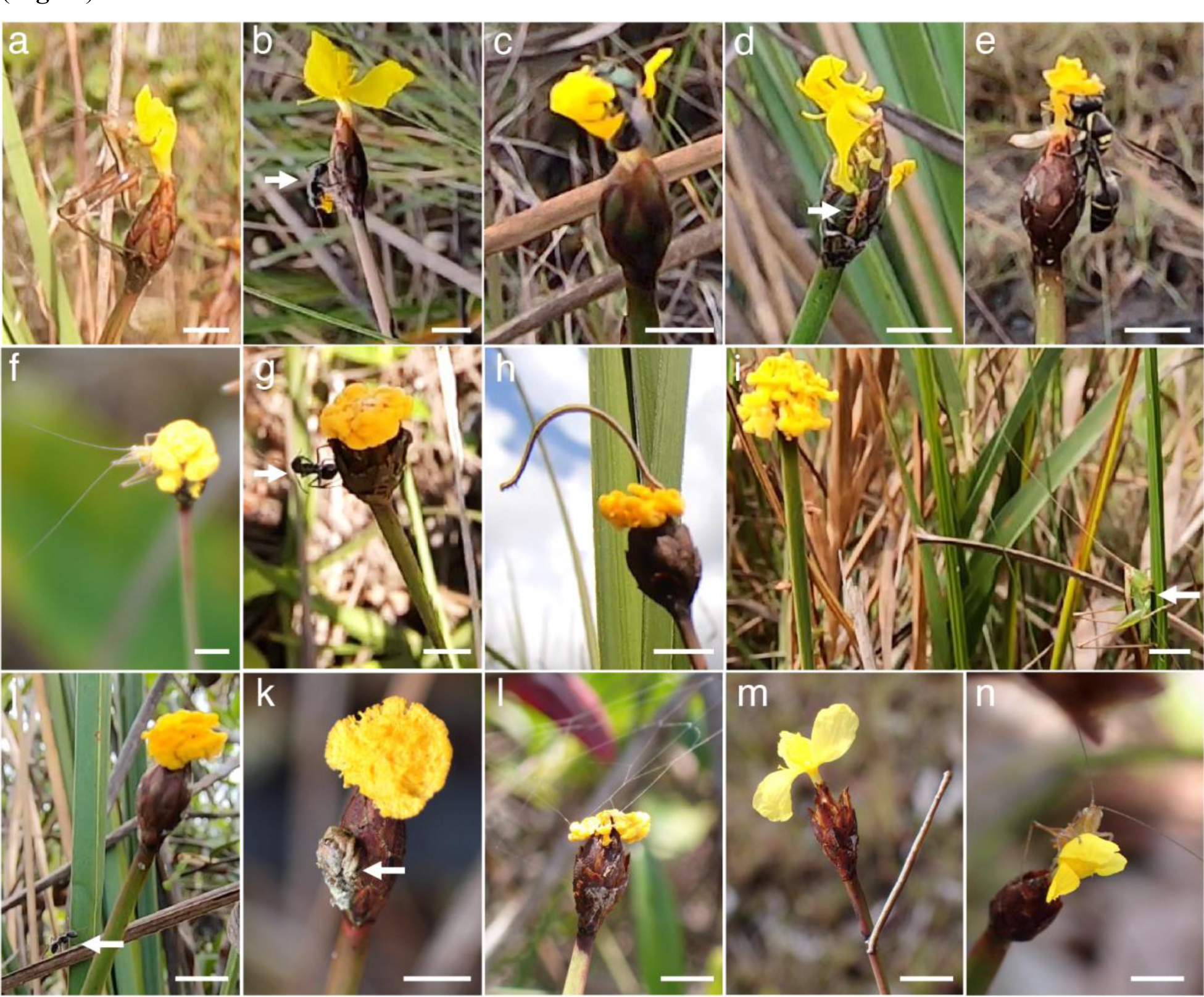
*Xyris surinamensis* flower and pseudoflower arthropod visitors. **a–e.** Insects in contact with flowers of *X. surinamensis*. **a.** Conocephalini sp. chewing on petals. **b.** *Camponotus* sp. carrying petal. **c.** Anthophila. **d.** Formicidae. **e.** Vespidae. **g–h.** Insects in contact with *Fusarium xyrophilum* pseudoflowers of *X. surinamensis*. **f.** Conocephalini sp. chewing on pseudoflower. **g.** *Camponotus* sp. **h.** Geometridae sp. **i.** Conocephalini sp. on *Xyris* leaf near a pseudoflower. **j.** *Camponotus* sp. on dead *Xyris* leaf near a pseudoflower. **k.** Salticidae sp. on *Xyris* spike below pseudoflower structure. **l.** Spider web in contact with pseudoflower. **m.** Geometridae sp. attached to stem below inflorescence. **n.** Conocephalini sp. chewing on petals. Pictures A–J are still images from timelapse videos recorded with GoPro Hero7 and K–N are photographs taken with an Olympus Tough TG-6 while in the field. White arrows point to small arthropods for easier visualization. Scale bar = 10 mm

Many of the arthropod visits to *Xyris* tissues lasted for less than a minute (**Table 3**). Yet, several Formicidae interactions with the flowers lasted between 1 min and up to 12 min and 30 s (**Table 3**). An Apidae sp. also remained in contact with a flower for 1 min and 4 s (**Fig. 1c**). Additionally, a Geometridae sp. (**Fig. 1h**) and a Conocephalini sp. (**Fig. 1f**) interacted with pseudoflowers for 2 min 15 s and 5 min 2 s, respectively. The longest interactions were captured between *X. surinamensis* and Conocephalini spp. One of these interactions lasted for 27 min 30 s and on the other one it took a katydid 49 min to eat an entire flower (**Table 3**). Some insects also fed on pseudoflowers. Potential chewing damage was observed on pseudoflowers in the field (**Fig. 2e–h**), and our video footage captured a katydid chewing on a pseudoflower (**Table 3**, **Fig. 1f**), as well.

**FIG. 2.**
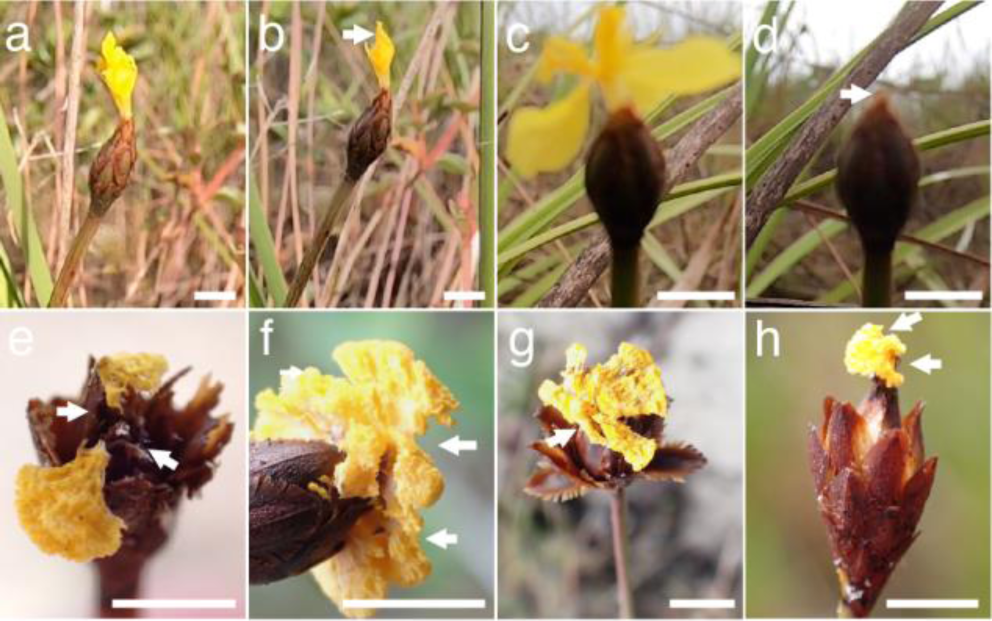
Evidence of insect feeding on *Xyris surinamensis* in Demerara-Mahaica, Guyana. **a–d.** Still images from timelapse recordings where Conocephalidae spp. were observed chewing on flowers. **a.** Flower right before arrival of Conocephalini sp. at site 1. **b.** Same flower after 8 min of Conocephalini sp. chewing on it. **c.** Flower before arrival of Conocephalini sp. at site 2. **d.** Same flower after 49 min of Conocephalini sp. chewing on it and eating all the floral tissues. **e–h.** Potential evidence of insect feeding occurring on pseudoflower tissues. White arrows point to areas where chewing has occurred. Scale bar = 10 mm

Fourteen other instances of insects were recorded visiting *Xyris* tissues in the periphery of the intended flower or pseudoflower being observed. In most cases, these were insects previously observed in contact with flowers and/or pseudoflowers that were in the frame of the recording but not come in direct contact with these tissues, such as Conocephalini spp. (*e.g.,* **Fig. 1i**) and several *Camponotus* spp. Other Formicidae and Acrididae sp. came in contact with *Xyris* leaves (*e.g.,* **Fig. 1j**).

### Molecular detection of *F. xyrophilum* on arthropod visitors

Eleven insects representing six families and one orb weaver spider (Araneidae), were hand-collected while in contact with *X. surinamensis* flowers or *F. xyrophilum* pseudoflowers in the field (**Fig. 3**). We collected one specimen from each of the families Chrysomelidae, Formicidae, and Geometridae. Eight of the specimens captured were orthopterans: five Conocephalini (Tettigoniidae), two Acrididae, and one Tetrigidae (**Table 3**). The 12 arthropods collected were tested using IGS rDNA PCR primer pair targeting *F. xyrophilum*. Amplicons of expected size (i.e., ∼173 bp) were observed for a Conocephalini (Tettigoniidae; **Fig. 3i**) and a Tetrigidae (**Fig. 3l**) collected while in contact with pseudoflowers, and a *Leptysma* sp. (Acrididae; **Fig. 3k**) which was captured during a visit to a *Xyris* flower (**Table 4**). Sanger sequences of these three amplicons (OQ121925– OQ121927) confirmed their identity as *F. xyrophilum*.

**FIG. 3.**
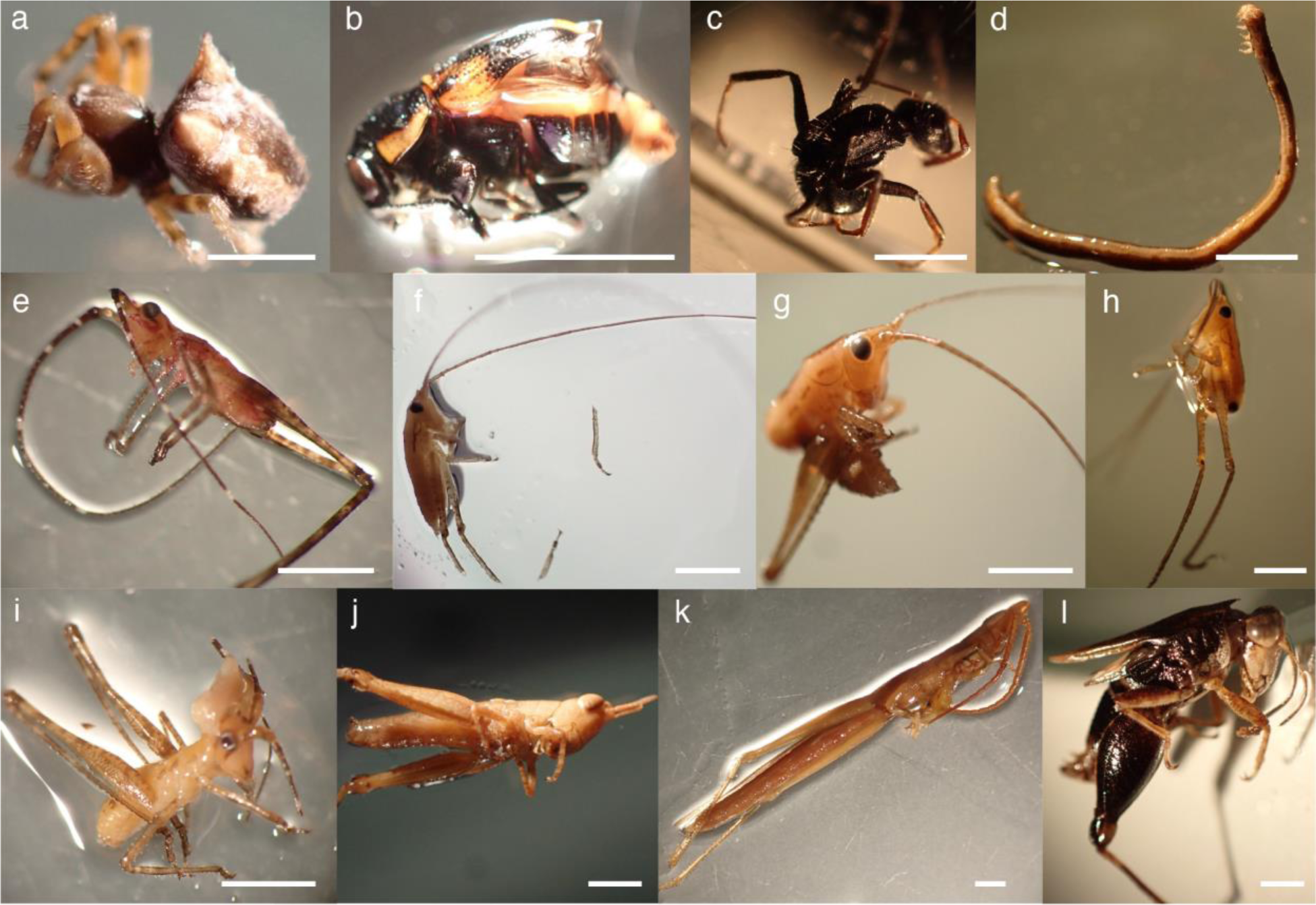
Macroscopic pictures before DNA extraction of arthropods collected by hand while in contact with *Xyris surinamensis* flower or pseudoflower tissues at Site 1, Demerara-Mahaica, Guyana. **a–c.** Arthropods found in contact with flowers. **a.** Araneidae (insect 27). **b.** Chrysomelidae (insect 26). **c.** *Camponotus* sp. (insect 31). **d.** Geometridae found attached to the stem right below the spike (insect 35). **e–h.** Conocephalinae (Tettigoniidae) found in contact with flowers (insects 30, 32, 33, 34; respectively). **i.** Conocephalinae (Tettigoniidae) found in contact with a pseudoflower (insect 25). **j.** Acrididae found on flower (insect 24). **k.** *Leptysma* (Acrididae) found on flower (insect 29). **l.** Tetrigidae collected while in contact with a pseudoflower (insect 28). **a–b.** Scale bar = 2.5 mm. **c–l.** Scale bar = 5 mm.

**Table 4.**
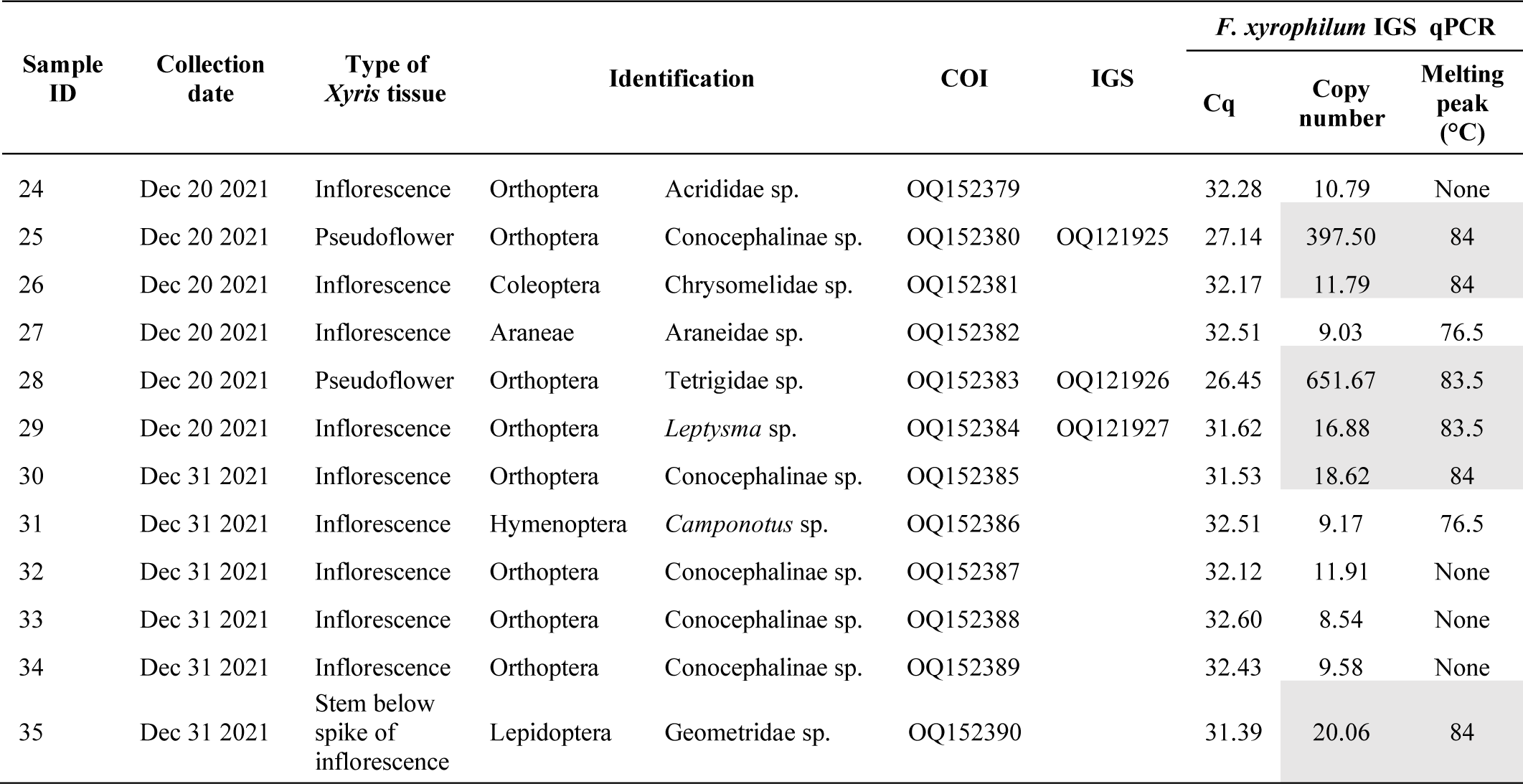
Presence of *F. xyrophilum* and identification of arthopods collected while visiting flowers and pseudoflowers on *Xyris surinamensis* at Site 1, Demerara-Mahaica, Guyana. Only results highlighted in grey are considered to be the correct IGS amplicon for qPCR

A qPCR of the 12 aforementioned arthropod samples using the IGS rDNA primers was also conducted to quantify the presence of *F. xyrophilum* on these insects. The serial dilutions of genomic *F. xyrophilum* DNA used for the curve of Cq vs. log number of amplicon molecules initially present demonstrated a reaction efficiency of 102.38% with an R^2^ of 0.991 (**Fig. 4a**), which meets the guidelines set out in Bustin et al. (2009). We were able to reliably amplify the concentration of the lowest standard containing ∼12 gene copies in the PCR tube at a Cq below 35. This sets the sensitivity of our assay towards *F. xyrophilum* IGS. Potential off-target amplification was observed in melting curves following qPCR reactions showing a peak at 76.5 °C (**Fig. 4b**), but this peak reached the cycle threshold (Ct value) in only one sample (insect 34) and the no template control (NTC). A subsequent gel electrophoresis revealed that for these samples with high 76.5 °C melting peaks only a band <50 bp was observed, that we suspect to represent primer dimer (data not shown), especially given the presence of this band in the NTC. Two samples (identifier 25 and 28; **Table 4**) presented especially high initial copy numbers, confirming the results from conventional PCR. However, several other insect samples (identifiers 26, 30, 35; **Table 4**) had detectable amounts of *F. xyrophilum* DNA that were not detected with conventional PCR and gel imaging. Sanger sequencing of selected amplicons followed by BLAST analysis showed, respectively, a 122/122 bp (100%) and 121/121 bp (100%) match to isolates SEr (MT919914) and NRRL 62710 (MT919908) of *F. xyrophilum*.

**FIG. 4.**
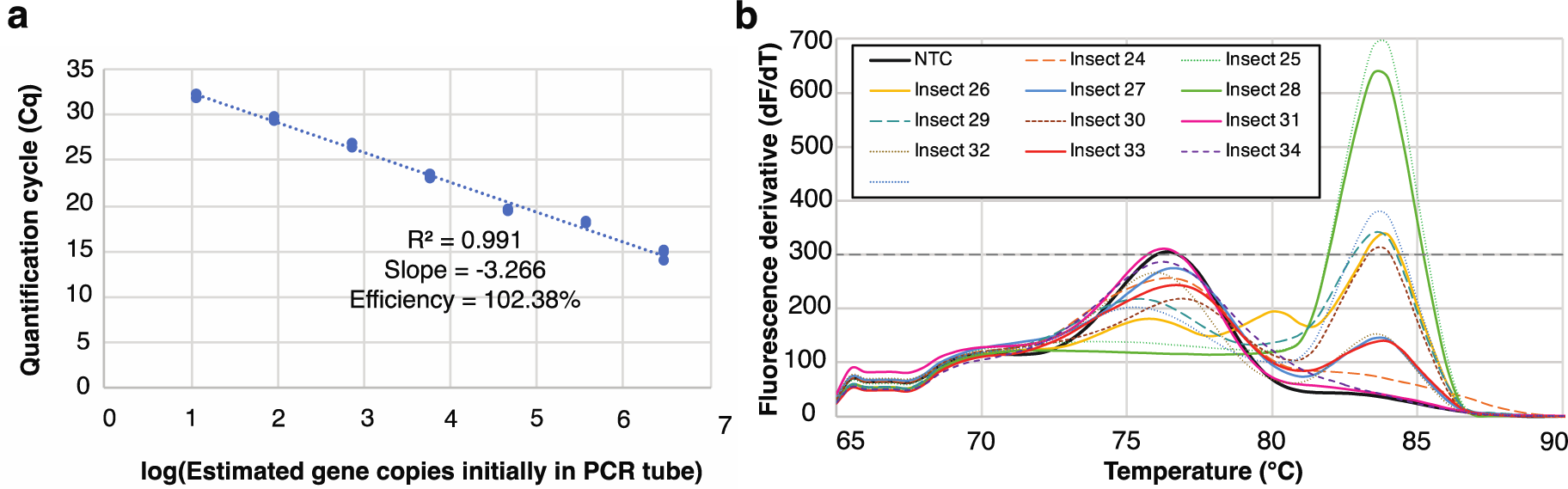
Calibration and melting curves for the IGS qPCR using primers IGS-1f and IGS-1r from Laraba et al. 2020. **a.** Calibration curve generated from a set of eight-fold standard dilutions of the IGS PCR product. **b.** Melt curves of qPCR products resulting from genomic DNA extracted from 12 insects that were hand collected from *Xyris surinamensis* tissues (flowers or pseudoflowers produced by *Fusarium xyrophilum*). Dotted line represents the cycle threshold (Ct value). NTC = negative control

### Volatile organic compounds emitted by *X*

surinamensis *flowers, and* Fusarium xyrophilum *pseudoflowers and cultures. Xyris surinamensis* flowers and *F. xyrophilum* pseudoflowers emitted (*Z*)-3-hexen-1-ol and 1-hexanol. *Fusarium xyrophilum* pseudoflowers, but not *X. surinamensis* flowers, emitted a sesquiterpene compound that eluted at 22.03 min, tentatively identified as α-gurjunene using the NIST14 library (**Fig. 5**). This sesquiterpene was also detected in the head space of *F. xyrophilum* PDA cultures (**Fig. 6**), but eluted at 29.55 min, presumably because a different method (i.e., SPME fiber) was used for volatile detection from fungal cultures. Interestingly, α-gurjunene was not present in the blend VOCs emitted by PDA cultures of *F. verticillioides* (NRRL 20956, FRC-M3125) and *F. thapsinum* (NRRL 22048, FRC-M6562) used for comparison. The mass spectra of α-gurjunene are provided in **Supplementary Material 2**. It is noteworthy that α-gurjunene was not detected in emissions from *X. surinamensis* flowers (**Fig. 5**).

**FIG. 5.**
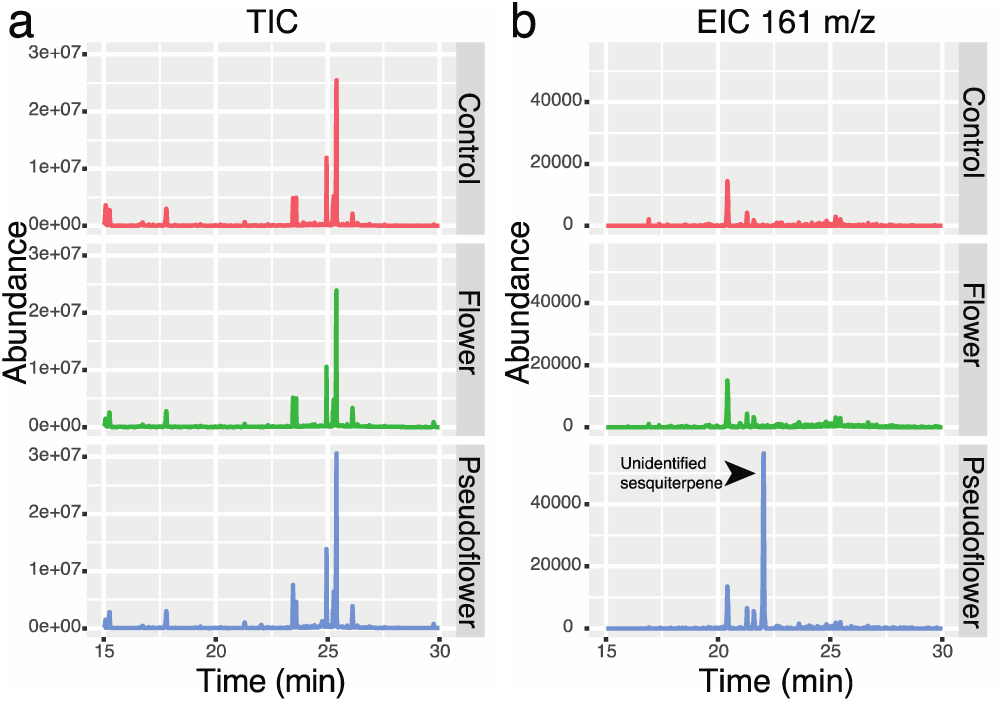
GC-MS spectrum of volatiles emitted from flowers or pseudoflowers of *Xyris surinamensis* in Guyana. **a.** Total ion chromatogram (TIC) **b.** Extracted ion chromatogram (EIC) for 161 m/z. Arrowhead points to sesquiterpene peak identified potentially as α-gurjunene

**FIG. 6.**
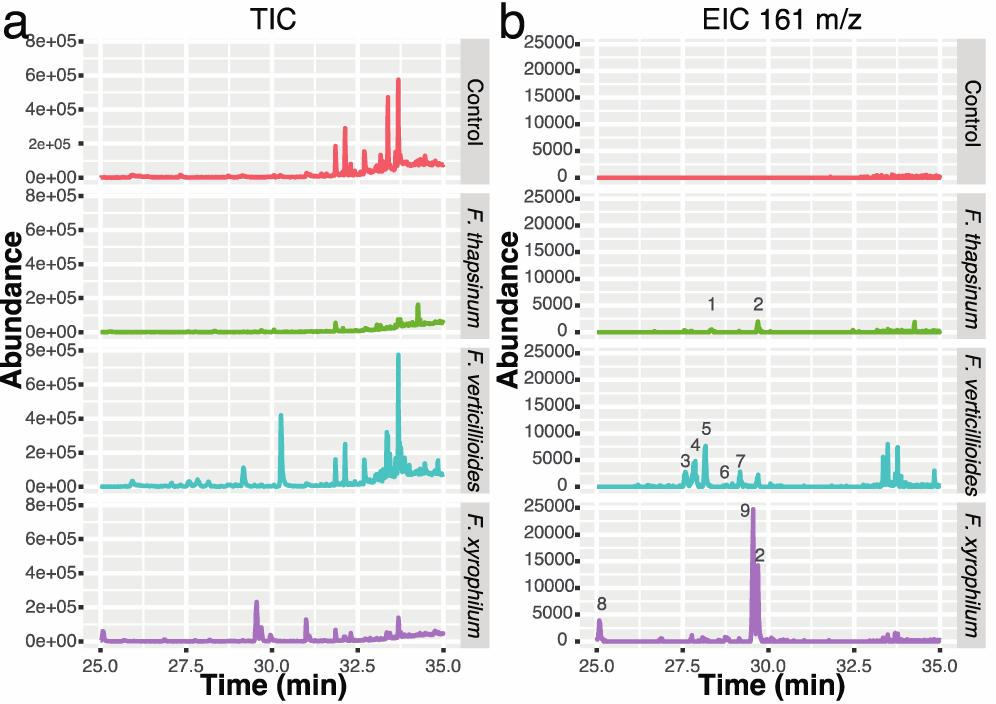
GC-MS spectrum of volatiles emitted from strains of *Fusarium thapsinum* (NRRL 22048, FRC-M6562), *F. verticillioides* (NRRL 20956, FRC-M3125), and *F. xyrophilum* (KOD596, NRRL 62721, FRC-M8921) grown on PDA. **a.** Total ion chromatogram (TIC) **b.** Extracted ion chromatogram (EIC) for 161 m/z. **1.** Dihydro-β-ionone. **2.** Germacrene D. **3.** 4H-1,4a-Methanonaphthalene, 1,5,6,7,8,8a-hexahydro-2,5,5,8a-tetramethyl-, (1.alpha.,4a.alpha.,8a.beta.)-. **4.** β-Funebrene. **5.** Cedrene. **6.** β-Copaene. **7.** Acoradiene. **8.** δ-Elemene. **9.** α- Gurjunene

We conducted antiSMASH and BLAST analyses to determine if any of the predicted terpene synthase genes in the *F. xyrophilum* genomes (GCA_008711575, GCA_008711615, and GCA_008711595) could be the α-gurjunene synthase gene. A limitation of this analysis was that, as far as we are aware, the α-gurjunene synthase gene has not yet been identified in any organism. Nevertheless, the antiSMASH and BLAST analyses indicated that the *F. xyrophilum* genome has 12 terpene synthase genes. The high level of amino acid sequence identity (>83%, **Supplementary Material 3**) of 10 of the *F. xyrophilum* terpene synthase genes to orthologs in *F. fujikuroi* suggests the *F. xyrophilum* enzymes have the same metabolic functions as the *F. fujikuroi* enzymes. The functions in terpenoid biosynthesis of nine of these genes are known, but none of the functions indicate that the corresponding enzyme could catalyze synthesis of α-gurjunene. We were unable to assign putative functions to three of the *F. xyrophilum* terpene synthase genes: genes KO596_3141, KO596_8005 and KO596_9253 in strain NRRL 62721 (**Supplementary Material 3**). KO596_3141 has a closely related *F. fujikuroi* ortholog whose function in terpene synthesis has not been determined. KO596_8005 and KO596_9253 do not have closely related orthologs in *F. fujikuroi* and BLAST analysis did not provide clues as to terpenes the corresponding enzymes synthesize. Thus, KO596_3141, KO596_8005 and KO596_9253 are candidate α- gurjunene synthase genes.

### *Xyris surinamensis* flower and pseudoflower UV fluorescence

Imaging *F. xyrophilum* pseudoflowers irradiated with UV light at a wavelength of 365 nm did not cause any fluorescence of the structures (**Fig. 7a**). By contrast, at the same treatment of *X. surinamensis* plants caused bright green fluorescence of pedicels of both excised flowers (**Fig. 7b**) and intact flowers in the field (**Fig. 7c**).

**FIG. 7.**
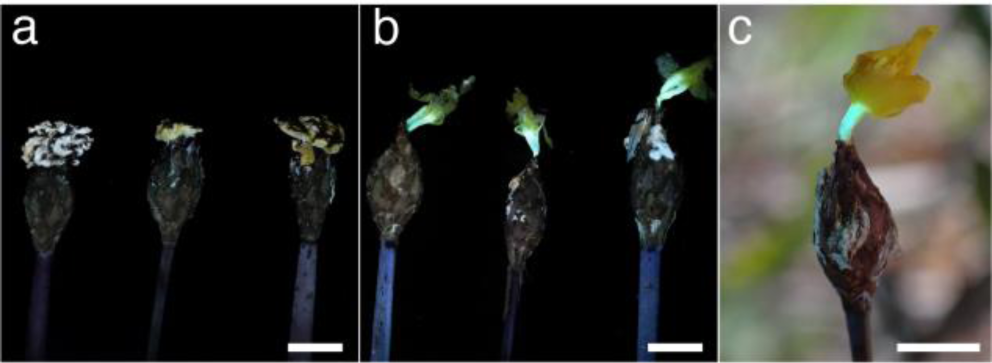
Fluorescence at 365 nm of *Xyris surinamensis* flowers and *Fusarium xyrophilum* pseudoflowers. **a–b.** Pictures taken approximately 6 h after removal from plant. **a.** Three *F. xyrophilum* pseudoflowers. **b.** Three *X. surinamensis* flowers. **c.** *Xyris surinamensis* flower in the field. Scale bar = 10 m

## DISCUSSION

Despite the lack of nectaries and ‘infrequent’ insect visitation previously reported for *Xyris* (Kral 1983), diverse arthropods were present in *Xyris surinamensis* populations in savanna areas of Guyana. Our study documents diverse insects in areas where *X. surinamensis* populations grow in Demerara-Mahaica Region of Guyana. Over 200 insects were captured with most individuals belonging to the orders Diptera and Hymenoptera. Dipterans from the families Syrphidae and Asilidae, as well as hymenopterans from Halictidae and Apidae, have been previously reported on *X. tennesseensis* populations from Albama and Georgia in the US (Boyd et al. 2011; Moffett and Boyd 2013), and *X. asperulla* and *X. tortulla* from Brazil (Freitas and Sazima 2006). However, herein, none of these families were observed on *X. surinamensis*. Dolichopodidae and Formicidae were the most common Dipteran and Hymenopteran families in our study, respectively. These differences may be due to bias in the collection methods used or perhaps Dolichopodidae and Formicidae may be more prevalent around *Xyris* populations in Guyana than those in temperate regions.

### Overlap in arthropods that visit *X. surinamensis* flowers and pseudoflowers

We evaluated the possibility that insect visitation to *Xyris* could be affected by the presence of *F. xyrophilum* pseudoflowers. It had been proposed that few insects infrequently visit *Xyris* species (Kral 1983). Follow-up studies showed a higher diversity of insect visitors, yet most of these studies were in North American *Xyris* species (Wall et al. 2002; Landry 2005; Freitas and Sazima 2006; Boyd et al. 2011; Moffett and Boyd 2013). First reports of arthropods in the orders Araneae, Coleoptera, Lepidoptera, and Orthoptera visiting *Xyris* spp. plants were recently made (Torres-Cruz et al. 2024); providing a more diverse view of the insects that visit these plants. Here we present the first assessment of insect visitors of *X. surinamensis* flowers and *F. xyrophilum* pseudoflowers.

Approximately 78 hours of video footage of insects visiting the flowers and pseudoflowers in the study area revealed a low number of insects in contact with *X. surinamensis* flowers (n =19) and pseudoflower tissues (n = 4). This number of observations is much lower than those noted in other fungal mimicry systems. For instance, in *B. vulgaris* infected by P*. arrhenatheri*, between 27–30 insect visitors were observed in 80 min and 148 in 360 min (Naef et al. 2002). On the other hand, in the *Mvc*-blueberry mimicry system, between ∼190–1450 bees and ∼60–250 flies visited true flowers but lower visitation rates (3–24 per hour) to *Mvc* infected leaves were documented (McArt et al. 2016). Despite the low number of insect visitors in this system, we provide the first report of Vespidae, Formicidae, and Acrididae visiting *Xyris* flowers and Salticidae on a pseudoflower. Salticidae have previously been observed to visit *Puccinia arrhenatheri* pseudoflowers on *Berberis vulgaris* (Naef et al. 2002). By contrast, the presence of Conocephalini spp. (katydids) is specific to this putative *F. xyrophilum*-*Xyris* mimicry system, as they have not been observed to visit other known fungal mimicry systems.

### *Xyris surinamensis flowers* and *F*

*xyrophilum* pseudoflowers were discovered to share certain visitors, including species of Conocephalini, *Camponotus*, and Geometridae, suggesting potential vectoring of *F. xyrophilum* conidia between uninfected and infected *Xyris* plants. While most arthropod visits to *Xyris* tissues lasted for less than a minute, some lasted up to 49 min where insects (e.g., ants and katydids) fed on *Xyris* flowers or *F. xyrophilum* pseudoflowers. This is a well-known behavior for certain orthopterans that feed on pollen and petals as a supplementary source of nutrition (Tan et al. 2017) and has been reported in other *Xyris* species in Guyana (Torres-Cruz et al. 2024). *Xyris* can be a food source for insects, like syrphid flies that feed on the pollen of *X. tennesseensis* (Boyd et al. 2011), which is presumed food for *C. xyridella* larvae (Landry 2005). Here we expand florivory on *Xyris* to Formicidae spp. *Fusarium xyrophilum* produces aseptate microconidia on the pseudoflowers, but not the typical *Fusarium* multiseptated fusiform macroconidia (Laraba et al. 2020a). It is possible that these microconidia survive the insect gut and/or are carried on the insect’s body. Hence, these field observations of insects chewing on the pseudoflowers suggest they might be tricking herbivores rather than pollinators for dispersal.

### *Fusarium xyrophilum* is detected on the bodies of arthropods

Our detection of *F. xyrophilum* DNA on insect visitors of flowers and pseudoflowers supports their previous predicted role as vectors of *F. xyrophilum* propagules (Laraba et al. 2020b). Our field observations showed great diversity of insects in proximity of *X. surinamensis* plants. However, only a subset came in contact with *X. surinamensis* tissues. Therefore, we exclusively captured insects (n=11) and other arthropods (n=1) that visited host flowers or pseudoflowers right before they were captured. DNA of *F. xyrophilum* was detected on 25% of samples using a conventional IGS rDNA targeting PCR. Using qPCR, we also detected *F. xyrophilum* in three other insects (i.e., Chrysomelidae sp., Conocephalinae sp., and Geometridae sp.), which did not test positive for *F. xyrophilum* in the conventional PCR. This is potentially due to higher sensitivity of the qPCR in comparison to the conventional PCR. These qPCR data increase our detection of *F. xyrophilum* DNA on insects to 50%. As they carry the fungal DNA on or in their bodies these insects are, therefore, likely vectors of *F. xyrophilum.* Despite the low number of insects tested in our study, our results are comparable to those from the *Mvc*-blueberry system where *Mvc* DNA was detected in ∼ 33% of captured bees and flies, the potential vectors of *Mvc* spores (McArt et al. 2016).

### Emission of a sesquiterpene by *F. xyrophilum* pseudoflowers and cultures

Pseudoflower-inducing fungi lure insects to infected plants through color but also using olfactory signals (Roy 1994; Naef et al. 2002; McArt et al. 2016). The volatiles produced by *F. xyrophilum* pseudoflowers were compared to those emitted by *X. surinamensis* flowers, *in situ* in Guyana to determine whether the pseudoflowers are mimicking the olfactory cues of the host flowers. *Xyris surinamensis* flowers emitted mainly two compounds, (Z)-3-hexen-1-ol and 1-hexanol, known green leaf volatiles likely formed after the flowers were excised (Ameye et al. 2018). This suggests that *X. surinamensis* flowers produce few, if any, constitutive volatiles. Conducting a more controlled collection under laboratory conditions in the future would be beneficial to further explore these results. Interestingly, pseudoflowers and *F. xyrophilum* cultures emitted a sesquiterpene compound tentatively identified as α- gurjunene. Our analysis of *F. xyrophilum* whole genome sequences detected three terpene synthase genes that are candidate α-gurjunene synthase genes. That is, the genome sequences included 12 terpene synthase genes, and the functions of nine of these genes is known based on closely functional analyses of closely related orthologs in other *Fusarium* species, whereas the functions of closely related orthologs of the three candidate α-gurjunene synthase genes have not been determined. α-gurjunene was not detected from *X. surinamensis* flowers or cultures of *F. verticillioides* and *F. thapsinum*, which are close relatives of *F. xyrophilum*. Despite not assessing the volatile profile of all members of the FFSC, these data suggest that α-gurjunene might be a *F. xyrophilum*-specific volatile with potential for involvement in insect attraction by pseudoflowers.

### UV fluorescence of *Xyris surinamensis* flowers

Insect pollinators are sensitive to the UV range of the electromagnetic light spectrum in addition to the visible spectrum (Briscoe and Chittka 2001). UV photoreceptors aid floral visitors in locating individual flowers that provide specific UV patterns, differing from other plants within the same community (Johnson and Andersson 2002). Our examination of *X. surinamensis* flowers and *F. xyrophilum* pseudoflowers revealed flower pedicels were UV fluorescent, but not the pseudoflowers. Blue UV-induced fluorescence emissions at 366 nm have been described on the floral parts, fruits, and seeds of several grasses (e.g., *Oryza sativa*, *Triticum aestivum*, *Zea mays*, *Sorghum bicolor*, *Ochlandra travancorica*, *Eleusine coracana*) and are suggested to be visual cues that attract pollinators to nectar (Thorp et al. 1975), petals (Gandía-Herrero et al. 2005), and pollen (Mori et al. 2018). On the other hand, it has been proposed that the fluorescence quantum efficiency of floral pigments is low (∼1%) and a fluorescence effect under natural conditions may be swamped by petal reflections (Iriel and Lagorio 2010). Therefore, fluorescence is currently regarded as ‘unimportant’ for visual signaling for pollinators (Van Der Kooi et al. 2019). Nevertheless, it is important to highlight that this has been evaluated from the perspective of UV fluorescence on the petal surface, which is superimposed on to the light reflected by the organism (Iriel and Lagorio 2010). However, in our observations *X. surinamensis* fluorescence was located on the pedicel, at the base of the flower, and not on the petals. The bright fluorescence of pedicels on *X. surinamensis* flowers may or may not be involved in insect attraction, and it should be evaluated with respect to the photoreceptor responses in the insect visitors’ eyes. Additionally, much of the knowledge of floral reflectance is based on studies of bees. Our studies of insect visitation to *Xyris* sp. and *X. surinamensis* in Guyana indicate the frequency of visits by bees is limited (Torres-Cruz et al. 2024). Nonetheless, it may be that these different cues act together to attract insect visitors to *Xyris* flowers and *F. xyrophilum* pseudoflowers. Whether or not, and to what extent, the UV reflectance and color of both flowers and pseudoflowers are visual signals involved in insect attraction remain to be determined.

## CONCLUSIONS

Here, we have expanded the list of *Xyris* insect visitors to include Vespidae, Formicidae, Salticidae, Acrididae, and Tetrigidae. Our results exemplify the need to study tropical regions, with an emphasis on remote and underexplored locations, to expand our knowledge of interactions in different ecosystems, often subject to sampling bias. Although we were unable to quantify differences in insect visitation to flowers and pseudoflowers, we have confirmed visitation to pseudoflowers by a diverse array of arthropods. Potential vectoring of *Xyris* pollen and *F. xyrophilum* conidia between plants is supported by 1) an overlap on the identity of insect visitors of *Xyris* flowers and *F. xyrophilum* pseudoflowers (Conocephalini, *Camponotus*, Geometridae), 2) observations of pollen on insects’ antennae and other parts of their bodies (Torres-Cruz et al.2024), and 3) evidence of *F. xyrophilum* DNA on insect bodies confirmed by IGS rDNA targeting PCR. It is also important to highlight that the interaction of *Xyris* and *F. xyrophilum* with other organisms could affect insect visitation. It has been suggested that microbial effects on pollen could influence pollen-eating animals, as they could plausibly affect pollen scent, nutrition, or physiology. Nevertheless, this topic has received little experimental attention (Vannette 2020). Additionally, we detected the emission of the sesquiterpene α-gurjunene on pseudoflowers and *F. xyrophilum* pure cultures but not by *Xyris* flowers, which could function as an insect attractant. Pseudoflowers have been shown to possess two pigments with fluorescence emission maxima in light ranges to which trichromatic insects are sensitive (Laraba et al. 2020b), and we have determined that the peduncle of *X. surinamensis* flowers fluoresces under UV light. These two visual cues, from flowers and pseudoflowers, paired with the different volatile emissions of pseudoflowers may work together to attract the high and diverse pool of insects we have observed visiting *Xyris*. Our study has also provided fundamental knowledge of insect interactions with *Xyris* flowers and *F. xyrophilum* pseudoflowers, including the first evidence for insect visitations, volatile production and fluorescence, and confirmation that insects disperse *F. xyrophilum*. Despite these advances in knowledge, much remains to be determined in the interaction of *Xyris* and *F. xyrophilum*. Our hypotheses for insect vectoring of *F. xyrophilum* and the involvement of insects in this potential mimicry system will need to be evaluated under extensive field experimentation.

## ACKNOWLEDGEMENTS

We thank Natalie Imirzian for advice on timelapse video logistics, Bitty Roy for input on methods for the insect visitation study, and Luciano and Francino Edmund, Patamona fungal parataxonomists, for their valuable help in field collection. We acknowledge the Guyana Environmental Protection Agency, the National Agricultural Research and Extension Institute, and USDA APHIS for permitting related to this project.

## STATEMENTS AND DECLARATIONS

### Funding

TJTC was supported by National Science Foundation grant DEB-1655980 and the Indigo Ag Phytobiomes Graduate Fellowship from the Penn State Microbiome Center. Field work by TJTC was partially funded by the American Philosophical Society Lewis and Clark Fund for Field Exploration and the Mycological Society of America Clark T. Rogerson Student Research Award; field work by MCA, LAR, AV,and JRJ was supported by National Science Foundation DEB-2127290 to MCA. TJTC thanks the Penn State University Department of Plant Pathology and Environmental Microbiology for the James and Marilyn Tammen and the Larry J. Jordan Endowments that partially supported costs for this project; as well as the Penn State Global Safety Office for the Wilderness First Aid Training Grant in preparation for field work in remote locations for this research project. DMG’s laboratory is supported by the Penn State Agricultural Experiment Station Project 4655. This work was supported in part by the U.S. Department of Agriculture, Agricultural Research Service.

### Competing Interests

The authors have no relevant financial or non-financial interests to disclose.

### Author Contributions

Terry Torres-Cruz developed the initial concept of the study, secured funding, led the fieldwork and laboratory analyses, analyzed the data, and wrote the first draft of the manuscript. Tristan M. Cofer advised on VOC experimental design, provided training in VOC sample collection, carried the VOC analysis, and interpreted VOC measurements. Laura M. Kaminsky provided training in qPCR technique and interpretation of results. Lauren A. Ré performed field work and analyzed video data. Jack R. Johnson and Alexis Vaughn supported field work and sample collection. James H. Tumlinson advised on VOC experimental design and provided funding for the VOC analysis. Imane Laraba advised on VOC sampling of *Fusarium* cultures. Robert H. Proctor and Hye-Seon Kim performed the antiSMASH analysis of terpene synthase genes in the *Fusarium* species. Terrence H. Bell advised on qPCR analysis. M. Catherine Aime supported field work logistics and funding for fieldwork assistants. Michael J. Skvarla advised on insect specimen collection design and supported insect identification. David M. Geiser participated on concept development, planning of the study, funding acquisition, and supervised the findings of this work. All authors commented on previous versions of this manuscript and approved the final manuscript.

### Conflicts of Interest

The authors have no relevant financial or non-financial interests to disclose.

### Data availability

All data supporting the findings of this study are available within the paper and its Supplementary Information files. Sequences were submitted to GenBank under accession numbers OQ121925–27 and OQ152379–90. Photography of plant, insect, and fungal observations at each collection site are available in iNaturalist under the project: “<u>Ecosystem profiles of *Xyris* Research Sites</u>”. Video clips of insect visitation observations are available on YouTube as playlist “*<u>Fusarium xyrophilum</u>* <u>– *Xyris surinamensis*</u> <u>insect visitation study</u>” (playlist ID: PL19pSjmfC9cSceih6nlMnaBQj57ITfvq8).

### Disclaimers

USDA is an equal opportunity provider and employer. Mention of trade names or commercial products in this publication is solely for the purpose of providing specific information and does not imply recommendation or endorsement by the U.S. Department of Agriculture.

## Notes

### Competing Interest Statement

The authors have declared no competing interest.

## REFERENCES

Ameye M, Allmann S, Verwaeren J, Smagghe G, Haesaert G, Schuurink RC, Audenaert K (2018) Green leaf volatile production by plants: a meta-analysis. New Phytologist 220:655–658. 10.1111/nph.14671

Batra LR, Batra SWT (1985) Floral mimicry induced by mummy-berry fungus exploits host’s pollinators as vectors. Science (1979) 228:1011–1013. 10.1126/science.228.4702.1011

Boyd RS, Teem A, Wall MA (2011) Floral biology of an Alabama population of the federally endangered plant, *Xyris tennesseensis* Kral (Xyridaceae). Castanea 76:255–265. 10.2179/11-006.1

Briscoe AD, Chittka L (2001) The evolution of color vision in insects. Annu Rev Entomol 46:471–510. 10.1146/annurev.ento.46.1.471

Folmer O, Black M, Hoeh W, Lutz R, Vrijenhoek R (1994) DNA primers for amplification of mitochondrial cytochrome c oxidase subunit I from diverse metazoan invertebrates. Mol Mar Biol Biotechnol 3:294–299. 10.1071/ZO9660275

Freeman S, Shtienberg D, Maymon M, Levin AG, Ploetz RC (2014) New insights into mango malformation disease epidemiology lead to a new integrated management strategy for subtropical environments. Plant Dis 98:1456–1466. 10.1094/PDIS-07-14-0679-FE

Freitas L, Sazima M (2006) Pollination biology in a tropical high-altitude grassland in Brazil: Interactions at the Community level. Ann Mo Bot Gard 93:465–516. 10.3417/0026-6493(2007)93[465:PBIATH]2.0.CO;2

Gandía-Herrero F, García-Carmona F, Escribano J (2005) Floral fluorescence effect. Nature 437:334. 10.1038/437334a

Iriel A, Lagorio MG (2010) Is the flower fluorescence relevant in biocommunication? Naturwissenschaften 97:915–924. 10.1007/s00114-010-0709-4

Johnson SD, Andersson S (2002) A simple field method for manipulating ultraviolet reflectance of flowers. Can J Botany 80:1325–1328. 10.1139/b02-116

Kaminsky LM, Bell TH (2022) Novel primers for quantification of *Priestia megaterium* populations in soil using qPCR. Appl Soil Ecol 180:104628. 10.1016/j.apsoil.2022.104628

Kral R (1988) The genus *Xyris* (Xyridaceae) in Venezuela and contiguous Northern South America. Source: Ann Mo Bot Gard 75:522–722. 10.2307/2399434

Kral R (1983) The Xyridaceae in the Southeastern United States. J Arnold Arboretum 64:421–429. https://www.jstor.org/stable/43782114

Kvas M, Marasas WFO, Wingfield BD, Wingfield MJ, Steenkamp ET (2009) Diversity and evolution of *Fusarium* species in the *Gibberella fujikuroi* complex. Fungal Divers 34:1–21.

Landry J-F (2005) Two new species of *Coleophora* from the New World, with record of a new hostplant family for Coleophorines (Lepidoptera:Coleophoridae:Coleophorinae). Holarctic Lepidoptera 10:9– 15

Laraba I, Kim HS, Proctor RH, Busman M, O’Donnell K, Felker FC, Aime MC, Koch RA, Wurdack KJ (2020a) *Fusarium xyrophilum*, sp. nov., a member of the *Fusarium fujikuroi* species complex recovered from pseudoflowers on yellow-eyed grass (*Xyris* spp.) from Guyana. Mycologia 112:39–51. 10.1080/00275514.2019.1668991

Laraba I, McCormick SP, Vaughan MM, Proctor RH, Busman M, Appell M, O’Donnell K, Felker FC, Aime MC, Wurdack KJ (2020b) Pseudoflowers produced by *Fusarium xyrophilum* on yellow-eyed grass (*Xyris* spp.) in Guyana: A novel floral mimicry system? Fungal Genet Biol 144:103466. 10.1016/j.fgb.2020.103466

Ma LJ, Geiser DM, Proctor RH, Rooney AP, O’Donnell K, Trail F, Gardiner DM, Manners JM, Kazan K (2013) Fusarium pathogenomics. Annu Rev Microbiol 67:399–416. 10.1146/annurev-micro-092412-155650

Marasas WFO, Ploetz RC, Wingfield MJ, Wingfield BD, Steenkamp ET (2006) Mango malformation disease and the associated *Fusarium* species. Phytopathology 96:667–672. 10.1094/PHYTO-96-0667

McArt SH, Miles TD, Rodriguez-Saona C, Schilder A, Adler LS, Grieshop MJ (2016) Floral scent mimicry and vector-pathogen associations in a pseudoflower-inducing plant pathogen system. PLoS One 11:e0165761. 10.1371/journal.pone.0165761

Moffett JM, Boyd RS (2013) Management of a population of the federally endangered *Xyris tennesseensis* (Tennessee Yellow-Eyed Grass). Castanea 78:198–212. 10.2179/12-034

Mori S, Fukui H, Oishi M, Sakuma M, Kawakami M, Tsukioka J, Goto K, Hirai N (2018) Biocommunication between plants and pollinating insects through fluorescence of pollen and anthers. J Chem Ecol 44:591–600. 10.1007/s10886-018-0958-9

Naef A, Roy BA, Kaiser R, Honegger R (2002) Insect-mediated reproduction of systemic infections by *Puccinia arrhenatheri* on *Berberis vulgaris*. New Phytologist 154:717–730. 10.1046/j.1469-8137.2002.00406.x

Ngugi HK, Scherm H (2006) Mimicry in plant-parasitic fungi. FEMS Microbiol Lett 257:171–176. 10.1111/j.1574-6968.2006.00168.x

O’Donnell K, Rooney AP, Proctor RH, Brown DW, McCormick SP, Ward TJ, Frandsen RJN, Lysøe E, Rehner SA, Aoki T, Robert VARG, Crous PW, Groenewald JZ, Kang S, Geiser DM (2013) Phylogenetic analyses of RPB1 and RPB2 support a middle Cretaceous origin for a clade comprising all agriculturally and medically important fusaria. Fungal Genet Biol 52:20–31. 10.1016/j.fgb.2012.12.004

Raguso RA, Roy BA (1998) “Floral” scent production by *Puccinia* rust fungi that mimic flowers. Mol Ecol 7:1127–1136. 10.1046/j.1365-294x.1998.00426.x

Roy BA (1994) The effects of pathogen-induced pseudoflowers and buttercups on each other’s insect visitation. Ecology 75:352–358. 10.2307/1939539

Tan MK, Artchawakom T, Abdul Wahab RBH, Lee C-Y, Belabut DM, Tan HTW (2017) Overlooked flower-visiting Orthoptera in Southeast Asia. J Orthoptera Res 26:143–153. 10.3897/jor.26.15021

Thorp RW, Briggs DL, Estes JR, Erickson EH (1975) Nectar fluorescence under ultraviolet irradiation. Science (1979) 189:476–478. 10.1126/science.189.4201.476

Torres-Cruz TJ, Ré L, Johnson J, Geiser DM, Skvarla MJ. 2024. Diversity of arthropods that visit *Xyris* spp. (Xyridaceae): New observations from Guyana. P Entomol Soc Wash 125(2):246–255. 10.4289/0013-8797.125.2.246

Van Der Kooi CJ, Dyer AG, Kevan PG, Lunau K (2019) Functional significance of the optical properties of flowers for visual signalling. Ann Bot 123:263–276. 10.1093/aob/mcy119

Vannette RL (2020) The floral microbiome: Plant, pollinator, and microbial perspectives. Annu Rev Ecol Evol Syst 51:363–386. 10.1146/annurev-ecolsys-011720

Wall MA, Teem AP, Boyd RS (2002) Floral manipulation by *Lasioglossum zephyrum* (Hymenoptera: Halictidae) ensures first access to floral rewards by initiating premature anthesis of *Xyris tennesseensis* (Xyridaceae) flowers. Florida Entomologist 85:290–291. 10.1653/0015

