## Supplementary Material 2 for "Insects visit *Fusarium xyrophilum* pseudoflowers on the host *Xyris surinamensis* (Xyridaceae) and carry fungal DNA on their bodies"

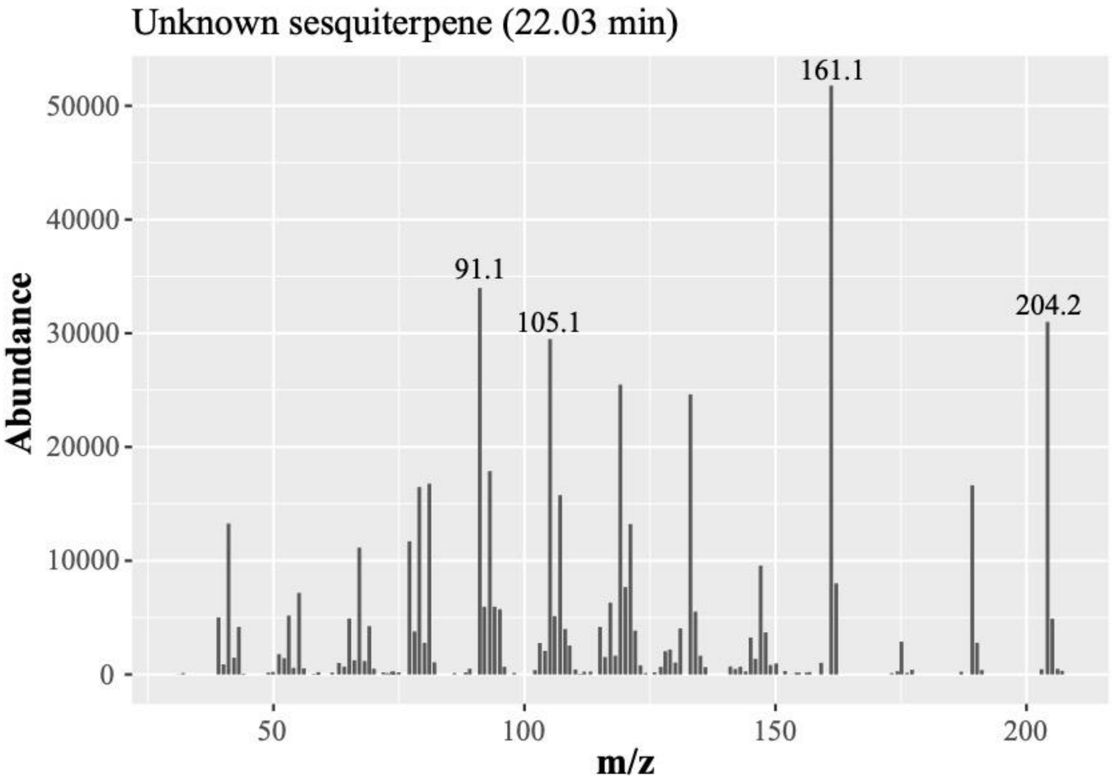


**Figure S1.** Reproduction of the mass spectrum (m/z) of the unknown sesquiterpene (22.03 min) detected on pseudoflowers of *Xyris surinamensis* in Guyana.

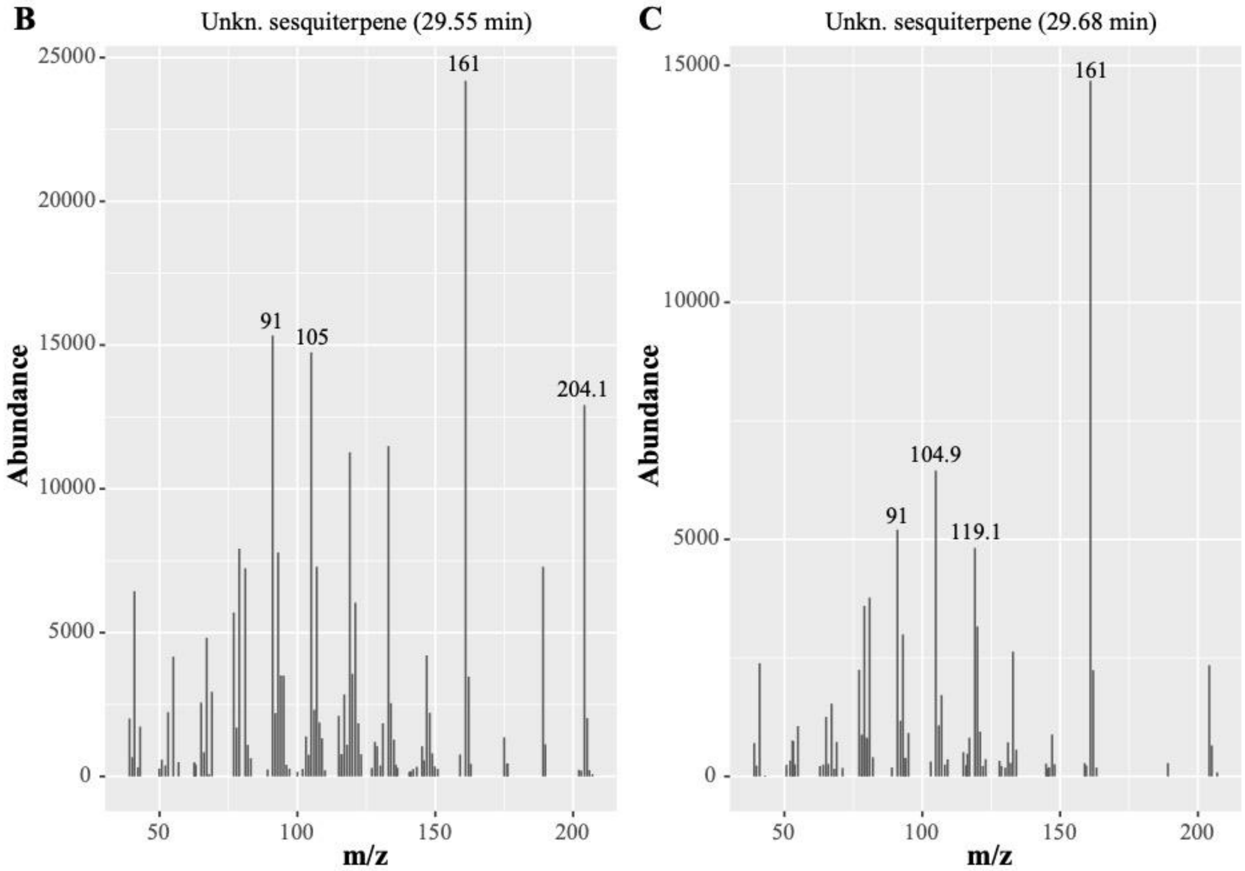


**Figure S2.** Reproduction of the mass spectrum **(**m/z) of unknown sesquiterpenes (29.55 and 29.68 min) detected on *Fusarium xyrophilum* (NRRL62721, FRC-M8921) grown on PDA.
